# A harmonized phantom MRI quality control framework identifies sources of longitudinal and multi-site variability

**DOI:** 10.64898/2026.09.29.754185

**Authors:** Balbino Yagüe Jiménez, Aref Kalantari, Declan Bolster, Elena Botto, Lorenzo Carnevale, Sabrina Doblas, Francesco Gammaraccio, Michael Gottschalk, Khan Hekmatyar, Jan Klohs, Nyoman D. Kurniawan, Xavier López-Gil, Bastian Maus, Edoardo Micotti, Federico Moro, Susanne Müeller, Lucia Perez Nieto, Chrystelle Po, Kelley M. Swanberg, Aurélien J. Trotier, Johan Van Audekerke, Annette Van der Toorn, Ignace Van Spilbeeck, Patricia Wenk, Thomas Basse-Lüsebrink, Daniele Bertoglio, Philipp Boehm-Sturm, Eike Budinger, João Miguel das Neves Duarte, Rick M. Dijkhuizen, Cornelius Faber, Gianluigi Forloni, Philippe Garteiser, William M. Holmes, Dario Livio Longo, Martin Meier, Emma Muñoz-Moreno, Emeline J. Ribot, Rui Vasco Simoes, Giovanna D. Ielacqua, Markus Aswendt

## Abstract

The shift towards open science in preclinical research requires high-quality, comparable data adhering to FAIR principles, yet rigorous quality assurance (QA) and quality control (QC) frameworks remain less established in preclinical magnetic resonance imaging (MRI) than in clinical imaging. To address this limitation, a multicenter study was conducted across 21 international laboratories using standardized commercial liquid phantoms for mouse and rat MRI setups and a harmonized acquisition protocol. Data processing was centralized using AIDAqc, an automated pipeline extracting quantitative metrics including signal-to-noise ratio (SNR), temporal SNR (tSNR), ghost-to-signal ratio (GSR), and motion-equivalent temporal instability, combined with five complementary outlier-detection algorithms. The evaluated datasets encompassed magnetic field strengths from 3.0 to 16.4 T and heterogeneous coil and acquisition configurations. In mouse phantom data, SNR and tSNR were significantly lower at 3 T than at 7 and 9.4 T, whereas artifact-related metrics did not differ significantly between field-strength groups. In rat phantom data, differences between field strengths were less pronounced. At 7 T, substantial dataset-specific differences were observed in anatomical and functional quality metrics, also among datasets using the similar coil configuration. Multicenter reference values for frequently represented 7 T configurations were established, including SNR/tSNR of 48 ± 3/48 ± 1 dB for mouse surface-coil datasets and 49 ± 3/44 ± 4 dB for rat 2 × 2 array-coil datasets. Longitudinal anatomical SNR showed generally low variability, whereas GSR was more variable across time. Automated outlier detection additionally identified transient acquisition instabilities that were not always apparent by visual inspection. Together, these findings show that field strength and nominal coil configuration alone do not adequately characterize MRI performance and support standardized, longitudinal phantom-based QC combined with automated analysis as a practical approach for improving the reliability, transparency, and interoperability of multicenter preclinical MRI data.

## Introduction

The recent paradigm shift towards open science in preclinical research has catalyzed a surge in multicenter studies, heavily underscored by the adoption of the FAIR (Findable, Accessible, Interoperable, and Reusable) data principles (Deruelle et al. 2020; Grandjean et al. 2023). However, the true value of shared imaging repositories hinges entirely on the underlying quality and comparability of the data. Quality assurance (QA) - the process ensuring quality in measurements and quality control (QC) - the measurement and detection process, are fundamental requirements for institutions generating imaging data. By systematically evaluating equipment using standardized QC protocols, QA processes including training, documentation, and monitoring ensure instrumental fidelity before any dataset is collected (Sreedher et al. 2021).

Preclinical magnetic resonance imaging (MRI) scanners are engineered for cutting-edge data acquisition, frequently utilizing ultra-high magnetic fields to deliver the spatial resolution required for small animal imaging (Marzola et al. 2003). However, differences in scanner hardware, acquisition configuration, and system performance can introduce technical variability that affects the reproducibility and comparability of quantitative MRI measurements, particularly in longitudinal and multicenter studies (Friedman et al. 2006; Jovicich et al. 2006). For preclinical MRI datasets to be genuinely interoperable and broadly reusable by the global scientific community, the instrumental conditions under which they were acquired must be rigorously reported and validated (Briggs et al. 2021). While MRI in the clinical setting has long operated under strict, standardized QA/QC mandates (Ihalainen et al. 2011; Chen et al. 2004), which are often enforced by regulatory agencies or as part of clinical trial contracts with industry partners, preclinical research has historically suffered from a lack of such rigorous frameworks (Osborne et al. 2017; Tavares et al. 2023). In the absence of standardized QA protocols and QC metrics across institutions, undetected hardware discrepancies may introduce confounding variables that compromise data integrity, rendering shared datasets fundamentally flawed for robust cross-validation.

Moreover, the urgent need for these frameworks is underscored by recent community surveys evaluating preclinical MRI practices (Osborne et al. 2017; Tavares et al. 2023). Despite the increasing use of small animal high field MRI scanners worldwide, institutional quality oversight remains alarmingly deficient. Crucially, survey results revealed that a staggering 63% of respondents either do not follow any QA/QC procedures or do not know of them (Tavares et al. 2023). Even when QC is attempted, it relies on a highly fragmented array of manufacturer-supplied, water-based, agar-based, or homemade phantoms to evaluate varying parameters like signal-to-noise ratio (SNR) or geometric accuracy.

To counteract these vulnerabilities, establishing a robust Standard Operating Procedure (SOP) should be paramount for any QA initiative. This initial benchmark acts as the ultimate reference point for longitudinal evaluations, playing a vital role in monitoring the long-term stability of the scanner. The continuous recording of these metrics is indispensable, not only allows researchers to detect performance drifts and anticipate potential hardware breakdowns, but it also establishes a verifiable audit trail needed to resolve inconsistencies in experimental data across different laboratories (Epistatou et al. 2020). Unified SOPs guarantee that measurements remain consistent regardless of the operator or the facility, establishing clear performance expectations and definitive action thresholds for every assessment.

To bridge this critical gap between the recognized need for standardized oversight and current laboratory practices, we present here a comprehensive multicenter study focused on QA in preclinical MRI. By deploying a unified QA framework across different institutions, this initiative aims to assess inter-laboratory reproducibility and continuously monitor the longitudinal stability of various scanner configurations. Ultimately, this collaborative effort seeks to lay the groundwork for acceptable QA guidelines, ensuring that preclinical imaging repositories can genuinely uphold the FAIR principles and deliver reliable, high-fidelity data to the scientific community.

## Material & Methods

### Study Design and Imaging Protocol

To evaluate the reproducibility and establish standardized QA/QC protocols in preclinical MRI, a multicenter international study was conducted. The progress was transparently reported^1^. We contacted 63 international MRI laboratories during 2023 to 2025 and received longitudinal phantom data from 21 sites. A longitudinal design was implemented to assess stability over time, with all centers required to acquire a minimum of four time points at weekly intervals. To standardize measurements and minimize sample bias across institutions, two types of standardized commercial phantoms (typically delivered with Bruker systems) were utilized (**Fig. 1A**). Both phantoms consisted of demineralized water, copper(II) sulfate, and sodium chloride, providing a filling appropriate for ^1^H NMR in the frequency range of 200–700 MHz.

1. Mouse phantom: A 15 ml Falcon tube (identified as MRI PHAN 1H IM M. HEAD) containing an aqueous solution of 1 g/L CuSO4⋅5H2O and 4.31 g/L NaCl.
2. Rat phantom: A 50 ml Falcon tube (identified as MRI PHAN 1H IM R. HEAD) containing an aqueous solution of 1 g/L CuSO4⋅5H2O and 3.6 g/L NaCl.

**Figure 1:**
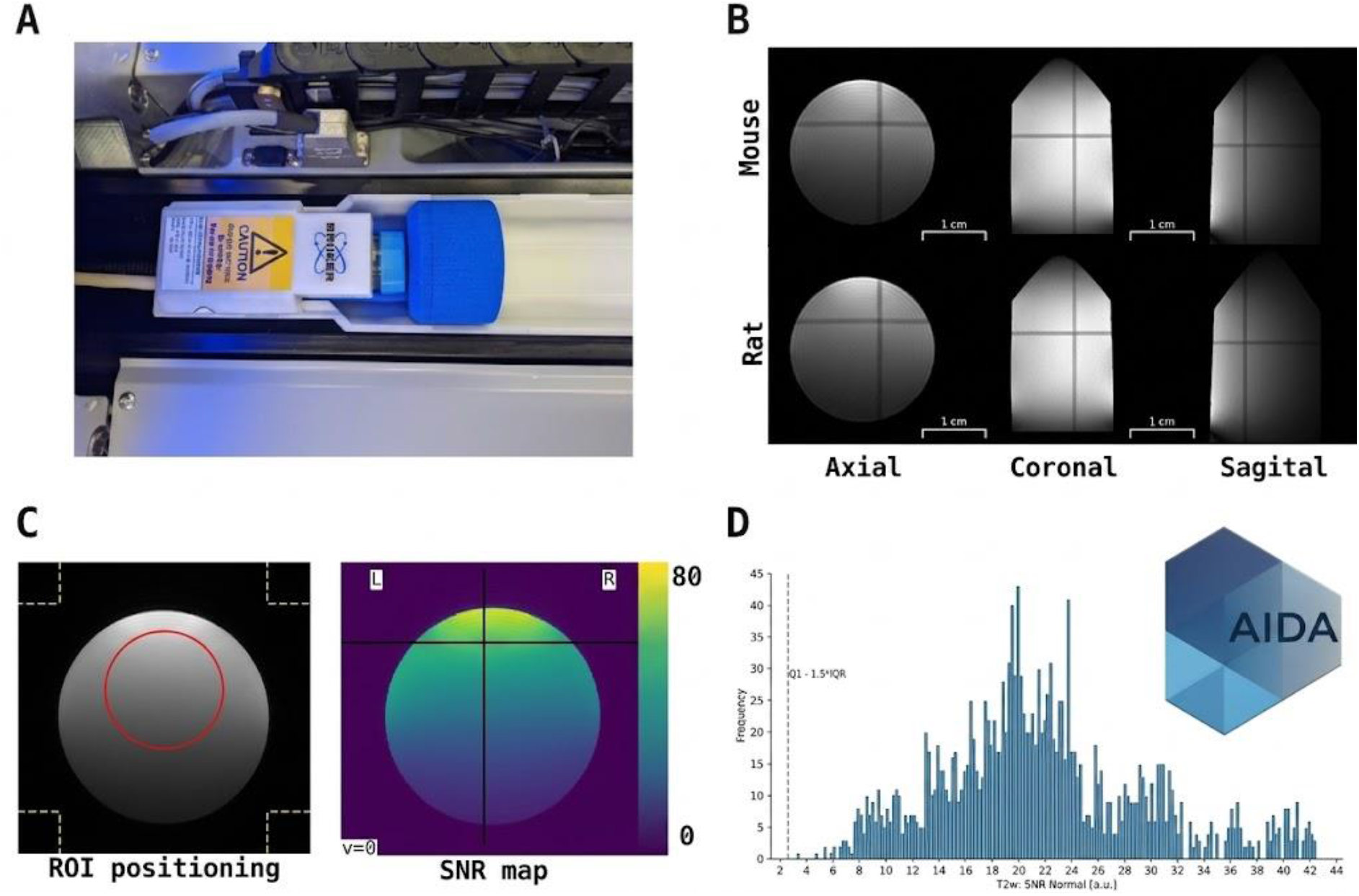
AIDAqc quality control workflow and standardized setup. (A) Representative experimental setup of a standardized rat 50 mL liquid MRI phantom. (B) Standardized positioning within the scanner for the mouse phantom (top row) and the rat phantom (bottom row). Dark lines indicate positions of the other oblique slices. Axial view (left), sagittal view (middle), coronal view (right). (C) Automated analysis interface showing the automatic region of interest (ROI) positioning (left) and a signal-to-noise ratio (SNR) map for rapid visual inspection (right). (D) Representative output histogram from the AIDAqc analysis displaying the frequency distribution of the normalized T2-weighted SNR.

To ensure technical reliability across different scanners, operators adhered to a harmonized acquisition protocol. The phantom was attached directly to the coil in a straight alignment, completely free of air bubbles. Positioning in the scanner’s isocenter was assessed using the localizer scan (**Fig. 1B**). To guarantee thermal equilibrium of the sample, the time elapsed from positioning the phantom inside the scanner bore to the start of the acquisition was kept strictly constant across all sessions (about 15 min). Prior to sequence execution, coil tuning and matching were mandatory, and the reference power of the coil was recorded for each session.

The sequence protocol encompassed both structural and functional simulated acquisitions: Anatomical Sequences (T1 and T2): T1WI and T2WI were acquired using Fast Spin Echo sequences. Both T1 (RARE) and T2 (turboRARE) sequences were configured with a RARE factor of 8 and fat suppression enabled. Functional Imaging Sequences: A single-shot FID gradient-echo echo-planar imaging (GE-EPI) sequence with T2-star contrast and 600 repetitions was used. The acquisition bandwidth was set to 300 kHz. Crucially, drift compensation, fat suppression, and AutoGhost correction modules were enabled (**Supplementary Material Table S1**).

### Data Management, Processing, and Quality Control

In accordance with current standards for transparency and reproducibility in neuroinformatics, all acquired raw data were curated and converted into the Brain Imaging Data Structure (BIDS) format (Gorgolewski et al. 2016). The raw data, the BIDS-compliant datasets, and the computed outputs were subsequently deposited and made publicly available using our previously established FAIR workflow for multimodal preclinical data (Kalantari et al. 2023). To eliminate inter-observer variability and centralize the QC process, all image processing and quantitative analyses were performed personally by a single investigator. The analysis was conducted using the AIDAqc platform, an ecosystem designed to address the reproducibility limitations in rodent functional MRI by providing standardized preprocessing, automated QC, and confound correction described in detail before (Kalantari et al. 2024). Briefly, the Python-based workflow comprises two main stages, initial parsing of raw MRI data and metadata followed by feature calculation and multivariate outlier detection. Extracted measures include SNR, tSNR, and motion-related metrics, whereas outlier detection combines 1) Interquartile Range (IQR): Effective for non-normal distributions based on data spread, 2) One-Class Support Vector Machine (ocSVM): Robust for capturing complex patterns in high-dimensional spaces, 3) Isolation Forest (IF): Efficiently isolates anomalies through decision tree construction, 4) Local Outlier Factor (LOF): Detects local density deviations, highly sensitive to local noise variations, 5) Elliptic Envelope (EE): Provides probabilistic detection based on robust covariance estimations. The outcomes of these five algorithms are aggregated using a majority voting system. Each scan receives an outlier severity score ranging from 0 (cleared by all) to 5 (flagged by all algorithms). This multialgorithm approach minimizes false positives and provides flexible thresholding for the multicenter evaluation (Kalantari et al. 2024). Reports are created as machine-readable CSV files and visual scan reports, including parameter distributions, pie charts for spatial resolution compliance, and automated middle-slice snapshots. These comprehensive outputs facilitate rapid visual inspection and are directly hosted alongside the BIDS-curated raw data in the GIN repository to guarantee full transparency of the QA/QC process (Kalantari et al. 2024).

### Quantitative Feature Calculations

AIDAqc feature calculation was modified to match the phantom geometry and include new features. SNR is a fundamental metric for evaluating hardware performance across different static magnetic fields and coil configurations. For structural sequences, AIDAqc derives the SNR (reported in dB) using two parallel automated approaches to avoid the bias of manual Region of Interest (ROI) placement. For this study, standard SNR calculation was used for anatomical images (T1 and T2 WI). This method automatically determines the Center of Intensity (COI) of the phantom image to place a spherical signal ROI (with its radius dynamically sized to 10% of the average image matrix dimensions) or adaptively an ellipsoid ROI for thin volumes representing the “true signal” (**Fig. 1C**). Simultaneously, eight cuboid ROIs are generated at the corners of the image volume. The voxel intensities from all eight corner regions are pooled, and one standard deviation across the pooled background voxels is used to sample the background noise. SNR was calculated according to equation (1):

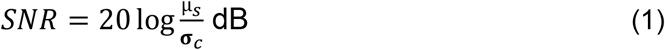

where μ_s_:is the mean signal intensity measured within the signal ROI, and σ_c_ is the standard deviation of the eight cuboids at the corners of the image volume.

Temporal Signal-to-Noise Ratio (tSNR): For functional GE-EPI sequences with successive volumes acquired during a single 4D imaging run, temporal signal stability during the GE-EPI acquisition was quantified using tSNR. The first 10 volumes were excluded when at least 10 volumes were available. An intensity-weighted center is determined from the temporal mean image across all acquired volumes, and a spherical or, for thin volumes, ellipsoidal ROI is placed at this center. tSNR was calculated according to equation (2):

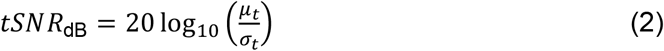

Where *μ_t_* is the temporal mean signal intensity of an individual voxel calculated across the entire time series and *σ_t_* is the temporal standard deviation of signal intensity for that same voxel across time.

The reported tSNR was the spatial mean of the voxel-wise dB-transformed tSNR values within the automatically defined central ROI. For both metrics, histogram reports are created for visual inspection (**Fig. 1D**)

Ghosting and Temporal Instability (Motion Equivalent): GE-EPI acquisitions are sensitive to phase inconsistencies that can result in Nyquist ghosting. AIDAqc detects ghosting using mutual information (MI) according to equation (3):

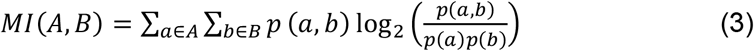

where *A*, *B* are the image slices being compared, *p*(*a*, *b*) is the joint probability distribution of signal intensities between images *A* and *B* and *p*(*a*), *p*(*b*) is the marginal probability distributions of signal intensities in images *A* and *B*, respectively.

For 4D datasets, a temporal mean image is first generated, whereas 3D datasets are analyzed directly. The geometrical middle axial slice is selected and circularly shifted successively along the second image-array axis. MI is calculated between the original slice and each shifted image. Peaks in the resulting MI profile are evaluated at successive dyadic offsets (½, ¼, ⅛ of the field of view, etc.). Peaks exceeding 25% of the maximum MI are classified as strong peaks, while all detected local maxima are additionally considered as weak peaks. Ghosting is flagged when at least one strong peak or more than two weak peaks coincide with the expected offsets. Although phantoms are stationary, temporal signal fluctuations can produce changes in image content resembling motion. AIDAqc therefore quantifies temporal instability using MI as a motion-equivalent metric according to equation (4):

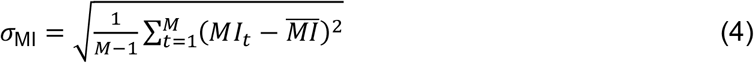

where *σ*_MI_ is the motion-equivalent temporal instability metric (standard deviation of *MI* values), *MI_t_* is the MI between the reference volume (*S*_1_) and the volume at time point *t* (*S_t_*), defined as *MI_t_* = *MI*(*S*_1_, *S_t_*), and 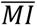 is the mean MI value across all retained time points

For 4D datasets containing at least 11 volumes, the first 10 volumes are excluded; otherwise, all volumes are retained. The axial slice with the highest mean signal, averaged across both in-plane spatial dimensions and the retained time series, is selected. The first retained volume of this slice is used as the reference, and MI is calculated between this reference and each subsequent retained volume. The standard deviation of the resulting MI values is reported as the motion-equivalent temporal-instability metric.

In addition to the Boolean MI-based ghosting flag, AIDAqc-Phantom calculates a continuous intensity-based ghost-to-signal ratio (GSR) for quantitative comparison of ghosting artifacts (Friedman and Glover 2006). For 4D datasets, the GSR is calculated from the temporal mean image across all acquired volumes; thus, unlike the temporal-instability calculation, the first 10 volumes are not excluded. The geometrical middle axial slice is selected, and an object mask is generated by thresholding positive-intensity voxels at the 70th percentile. If the resulting mask contains fewer than 10 voxels, the threshold is reduced to the 50th percentile; if fewer than 10 voxels remain, the GSR is reported as unavailable. The object mask is circularly shifted by half the field of view along the second image-array axis, corresponding to the assumed phase-encoding direction. The ghost-only region comprises voxels within the shifted mask that do not overlap the original object mask. Background intensity is estimated from voxels outside the original object mask. The GSR is calculated according to equation (5):

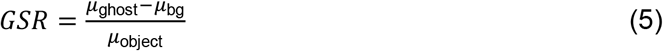

where *μ*_ghost_ is the mean intensity within the ghost-only region, *μ*_bg_ is the mean background intensity (estimated from voxels outside the original object mask), and *μ*_object_ is the mean intensity within the original object mask (voxels exceeding the 70th percentile threshold, or 50th percentile if < 10 voxels remain).

### Statistics

The statistical analyses were conducted using GraphPad Prism version 9.5.1 (GraphPad Software, San Diego, CA, USA), which was also used for data visualization. Mouse and rat phantom datasets were analyzed separately. For comparisons of MRI quality metrics (SNR, tSNR, motion-equivalent temporal instability, and ghost-to-signal ratio), the laboratory dataset was treated as the independent unit. When multiple acquisitions were available for the same dataset and time point, values were first averaged so that each time point contributed equally. For comparisons between field strengths and between datasets at 7 T, each dataset was represented by the mean across its available unique longitudinal time points. This dataset-level approach prevented datasets with a larger number of acquisitions from receiving greater weight in the statistical comparisons.

Temporal stability of anatomical MRI quality metrics was assessed using all available longitudinal T2_RARE acquisitions. SNR was quantified using the AIDAqc SNR_Normal metric and image ghosting using the ghost-to-signal ratio (GSR). For each dataset, temporal variability was quantified using the coefficient of variation (CV) according to equation (6):

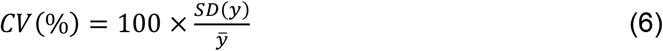

where *y* is the quality metric value evaluated across all available unique time points, *SD*(*y*) is the sample standard deviation of the quality metric *y*, and *y̅* is the arithmetic mean of the quality metric *y* across all unique time points.

Normalized drift therefore represents the estimated percentage change in the respective quality metric per measurement interval rather than cumulative change over the entire observation period. Normalized drift was calculated according to equation (7):

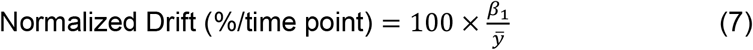

where *β*_1_ is the regression slope obtained from fitting a linear regression model of the quality metric *y* against measurement time points and *y̅* is the mean value of the quality metric *y* across the entire dataset.

Positive values indicate an increase and negative values a decrease over time. The standard error and 95% confidence interval of normalized drift were derived from the standard error of the regression slope, and the coefficient of determination (R^2^) was calculated to describe the proportion of temporal variance explained by the linear trend. For descriptive purposes, the mean and standard deviation of each quality metric across the available time points were additionally calculated for each dataset. Each laboratory dataset subsequently contributed one CV and one normalized-drift value per quality metric to the group-level temporal-stability analysis.

Statistical outliers in dataset-level temporal-stability measures were identified using Grubbs’ test (alpha = 0.05) and excluded from the corresponding group-level comparisons. Data distributions were assessed using the Shapiro–Wilk test for normality and the Brown–Forsythe test for homogeneity of variances. For comparisons of more than two groups, normally distributed data with homogeneous variances were analyzed using ordinary one-way ANOVA followed by Tukey’s multiple-comparisons test. When variances were unequal, Welch’s one-way ANOVA followed by Dunnett’s T3 multiple-comparisons test was used. Data that did not meet the assumptions for parametric testing were analyzed using the Kruskal–Wallis test followed by Dunn’s multiple-comparisons test. For comparisons between two groups, Welch’s unpaired t-test or the Mann–Whitney test was used, as appropriate. The statistical test applied to each analysis is specified in the corresponding figure legend, and complete statistical results are provided in the online supplementary material. Statistical significance was defined with significance levels indicated as * p < 0.05, ** p < 0.01, *** p < 0.001, **** p < 0.0001.

## Results

### Dataset overview and description

A standardized MRI protocol and processing pipeline was applied using mouse and rat phantoms from 21 independent laboratories (**Fig. 2**). The dataset captures the technological variability of hardware available across the community, i.e., participating centers utilized multiple radiofrequency (RF) coil configurations (**Supplementary Table S2**). The complete dataset included measurements from 3 to 16.4 T. There was only one 16.4 T dataset available and therefore this dataset was reported descriptively and excluded from statistical comparisons between field strengths. The longitudinal acquisitions of these datasets varied, although the majority of datasets consisted of the minimum requested 4 time consecutive time points, and a maximum of 12 acquisitions (**Fig. 2A**). MRI data were acquired all in Bruker systems, with a broad range of hardware in terms of field strength and coil arrangement (**Fig. 2B-C**). Most studies were conducted at 7 T for both rat (n = 12) and mouse (n = 11), followed by 9.4 T (n = 4 rat, n = 3 mouse), 3 T (n = 1 in rat, n = 2 mouse phantom) datasets and a 16.4 T mouse dataset (n = 1). Hardware setups varied further in terms of coil configuration across species. While mouse datasets were acquired mainly using single surface coils (n = 10), rat studies utilized a more heterogeneous arrangement, primarily favouring 2×2 phased-array surface coils (2×2 arrays, n = 7) followed by other types of coils (n = 6) (helmholtz + surface coil, different volume coils, 2×2 rat brain array cryoprobe and 3 x 1 array) (**Fig. 2C**). The datasets further varied substantially in terms of spatial resolution and voxel dimensions (**Fig. 2D-E**). Whereas rat imaging protocols generally clustered around a few specific voxel sizes, mouse datasets exhibited a high degree of fragmentation. Mouse protocols employed a wide multiplicity of voxel volumes (0.0118, 0.036, 0.003, 0.02, 0.2, 0.144, 0.08, 0.176, and 0.33 mm^3^).

**Figure 2:**
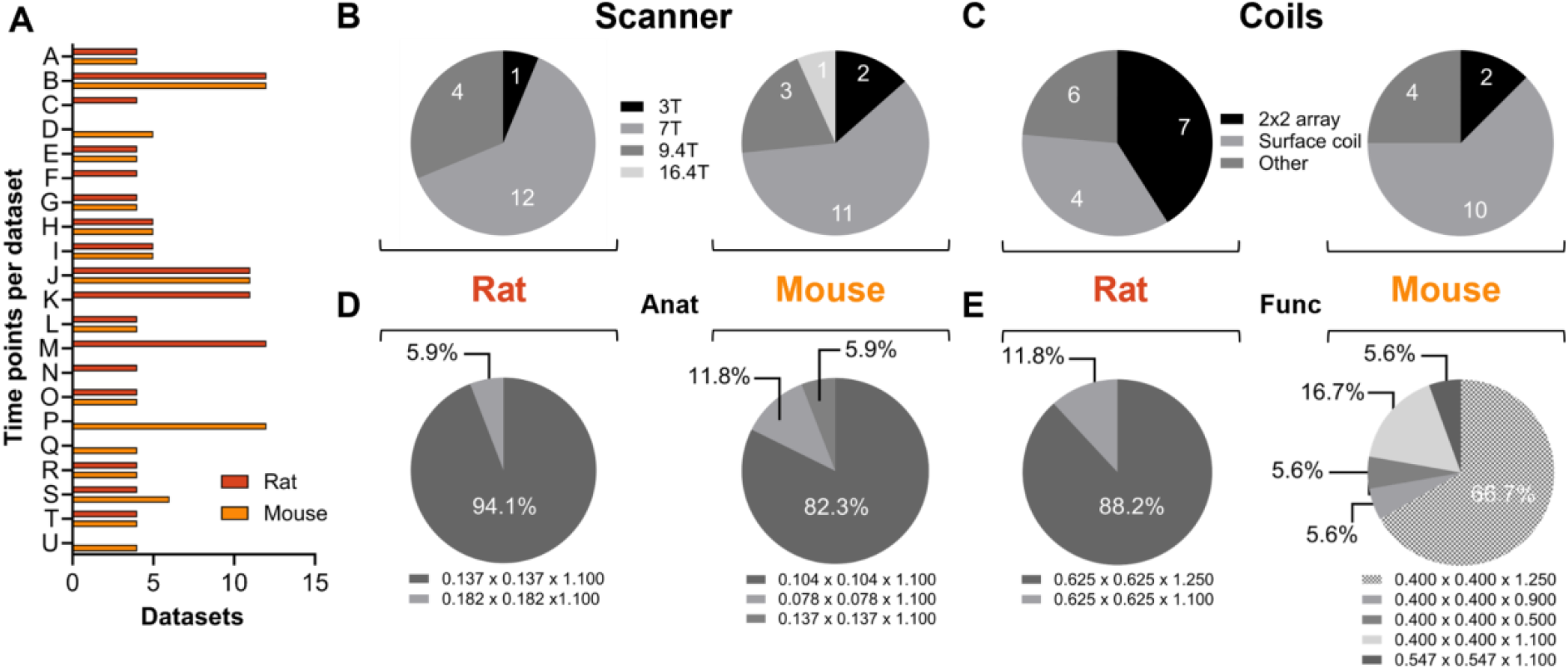
Phantom data acquisition workflow and quantitative comparison of resulting datasets. (A) Number of available QC time points for each site (anonymized as A–U), separated by mouse and rat phantoms. (B) Distribution of scanner field strengths across rat and mouse datasets, including 3, 7, 9.4, and 16.4 T systems. Numbers indicate the number of datasets for each field strength. (C) Distribution of receive coil configurations across rat and mouse datasets, grouped as 2×2 array, surface coil, or other coils. (D–E) Distribution of spatial resolutions (mm) and slice thickness (mm) for anatomical - T2-weighted (D) and spatial resolutions in voxel dimensions (mm) for functional - GE-EPI (E) MRI sequences, shown separately for rat and mouse phantoms. Pie charts indicate the relative contribution of the different voxel sizes and illustrate the heterogeneity of acquisition parameters across participating sites and scanner configurations.

### Global SNR and tSNR performance

We first compared MRI quality metrics (SNR, tSNR, motion-equivalent temporal instability, and ghosting as ghost-to-signal ratio) across magnetic field-strength groups for both rat and mouse phantoms (**Fig. 3**). In the rat phantom datasets (**Fig. 3A**), no significant differences in SNR were observed across the evaluated field strengths. tSNR was higher at 9.4 T than at 7 T, although the difference did not reach statistical significance (p = 0.0553). Artifact analysis in the rat phantom revealed no significant differences for either motion or ghosting. The single rat 3 T dataset is shown descriptively but excluded from comparisons. For the mouse phantom acquisitions (**Fig. 3B**), SNR at 3 T was significantly lower than at both 7 T and 9.4 T (p < 0.05). Similarly, tSNR was significantly lower at 3 T than at 7 T and 9.4 T (p < 0.001). No significant differences between field-strength groups were observed for motion or ghosting (**Fig. 3B**). The data acquired at 16.4 T was excluded because it was provided from one site only.

**Figure 3:**
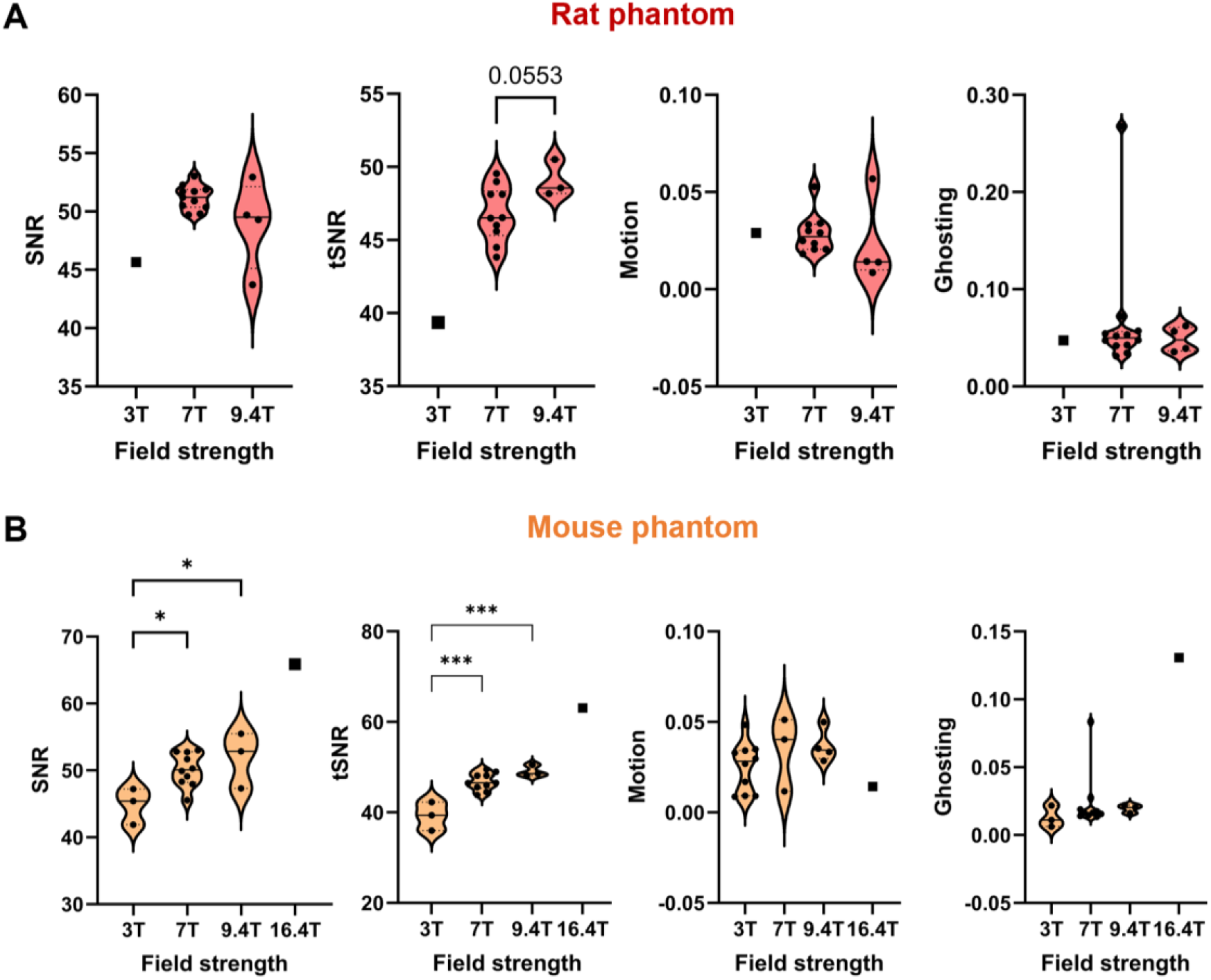
MRI quality metrics across magnetic field strengths in the rat and mouse phantom. Comparison of SNR, tSNR, motion, and ghosting for (A) rat and (B) mouse phantom datasets acquired at 3.0 T, 7.0 T, 9.4 T, and, for the mouse phantom, 16.4 T. Data are shown as violin plots with individual dataset values overlaid. Each symbol represents one independent laboratory dataset and corresponds to the mean of the respective quality metric across all available unique longitudinal time points. The number of available time points varied between datasets: rat phantom datasets comprised 11 time points at 3 T, 4–12 time points at 7 T, and 4–6 time points at 9.4 T; mouse phantom datasets comprised 11 time points at 3 T, 4–12 time points at 7 T, 4–6 time points at 9.4 T, and 12 time points at 16.4 T. When multiple acquisitions were available for the same dataset and time point, values were averaged before calculation of the dataset-level mean to ensure equal weighting of time points. Datasets include different RF coil and acquisition configurations within each field strength. Violin plots show the distribution of dataset-level mean values, with quartiles (dotted horizontal lines) and medians (solid horizontal lines). For the rat phantom, the single 3 T dataset was shown descriptively and excluded from inferential comparisons. For the mouse phantom, the single 16.4 T dataset was shown descriptively and excluded from inferential comparisons. Statistical comparisons were therefore performed between 7 and 9.4 T for rat datasets and between 3, 7, and 9.4 T for mouse datasets. Mouse metrics comparisons were performed using Ordinary one-way ANOVA for all parameters. Rat metric comparisons were evaluated using a Welch t-test for all parameters except ghosting, which was evaluated using a Mann-Whitney U test. Significant pairwise comparisons are indicated in the figure. *p ≤ 0.05, **p ≤ 0.001.

### Quantitative comparison of anatomical MRI quality across datasets and coil configurations at 7 T

Dataset-specific differences were observed in anatomical MRI quality measures at 7 T (**Fig. 4**). In the rat phantom, SNR was relatively stable over the four time points for most datasets, with differences in absolute SNR levels between datasets (**Fig. 4A**). These differences were limited, with only dataset O showing significantly higher SNR than dataset R (p = 0.028). More pronounced differences were observed for ghosting (**Fig. 4B**), both in absolute levels and longitudinal patterns. Dataset C showed particularly high ghosting and was significantly higher than eight of the ten other datasets (p= 0.010), with datasets G and H being the exceptions), including key comparisons highlighted in the figure such as A vs. C (p = 0.0095) and C vs. O (p = 0.0062). Dataset R showed comparatively low ghosting and was significantly lower than datasets A, C, E, H, I, and O (p = 0.006–0.045), with the illustrative comparison H vs. R (p = 0.0386). Despite the presence of multiple datasets acquired with surface and 2×2 array coils, neither SNR nor ghosting showed a consistent separation according to coil configuration. For the mouse phantom, SNR also varied between datasets (**Fig. 4C**); however, no pairwise differences remained significant after correction for multiple comparisons. In contrast, ghosting showed pronounced differences between individual datasets (**Fig. 4D**), with dataset G and U displaying particularly low values compared with several other datasets (p < 0.001).

**Figure 4:**
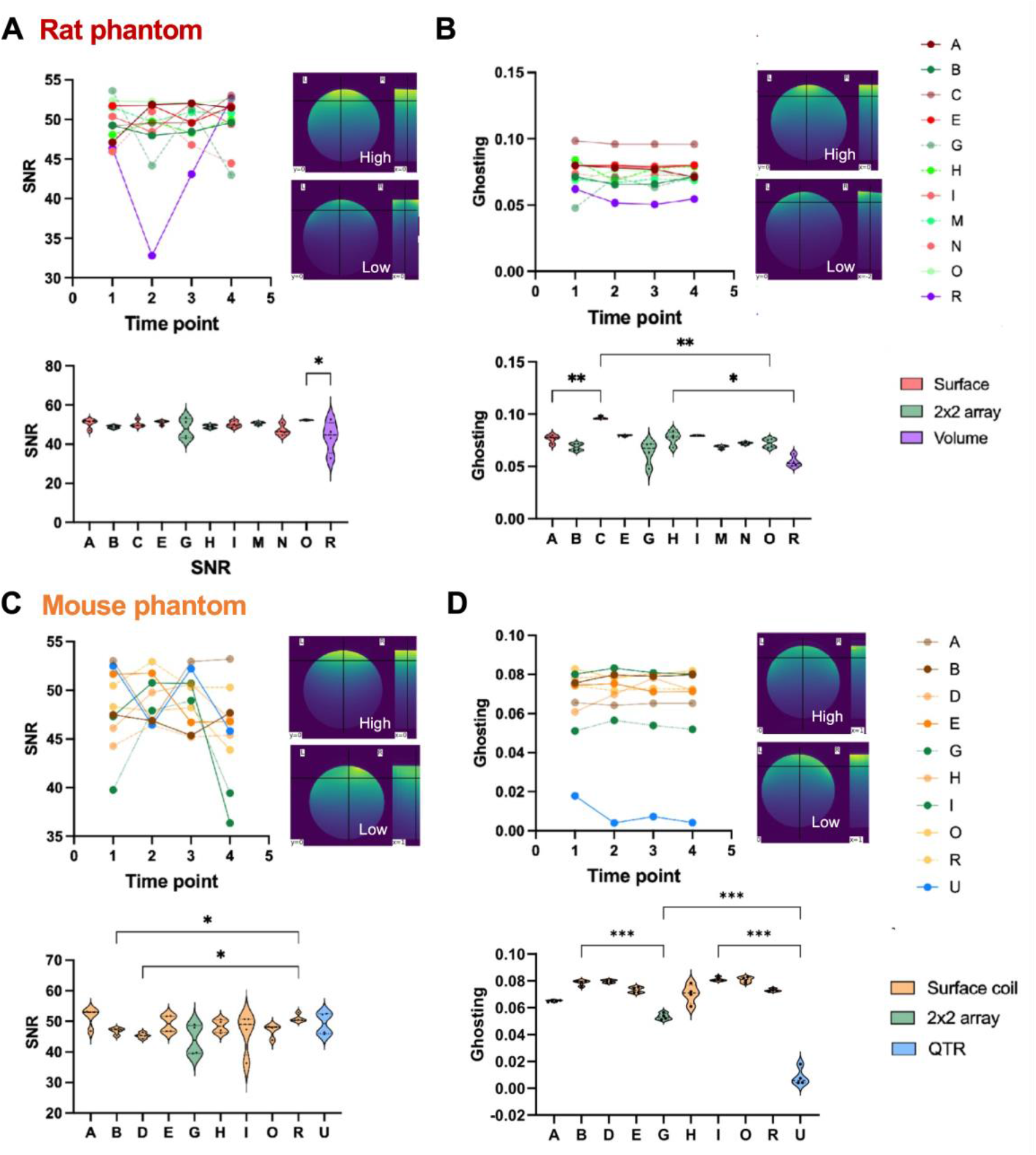
MRI quality measures for anatomical scans acquired with rat and mouse phantoms at 7 T. MRI quality measures for anatomical scans acquired with rat and mouse phantoms at 7 T. Quality control measures are shown separately for the rat (A–B) and mouse (C–D) phantoms, including SNR (A, C) and ghosting (B, D). Upper panels show individual datasets across four repeated time points; representative AIDAqc output images illustrate examples of high and low values for each quality metric (* indicates visible image artifacts). Lower panels compare the distributions of quality metric values across the four time points between datasets. Violin plots show the distribution of values, with quartiles (dotted horizontal lines) and the median (solid horizontal line). Statistics: ordinary one-way ANOVA followed by Tukey’s multiple-comparisons test (A, D), Brown–Forsythe and Welch ANOVA followed by Dunnett’s T3 multiple-comparisons test (B), and Kruskal–Wallis test followed by Dunn’s multiple- comparisons test (C). Selected significant pairwise comparisons are indicated in the figure. Detailed statistical results are available in the online data repository.

### Quantitative comparison of functional MRI quality across datasets and coil configurations at 7 T

Dataset-specific differences were also observed in functional MRI quality measures at 7 T (**Fig. 5**). In the rat phantom, tSNR was relatively consistent across the four repeated measurements within most datasets, whereas absolute tSNR levels differed between datasets (**Fig. 5A**). Dataset B showed the lowest mean tSNR (35.5) and was significantly lower than eight of the ten other datasets (p = 0.001–0.045), with datasets H and O being the exceptions. Rat phantom ghosting showed more pronounced dataset-specific patterns, both in absolute levels and across repeated measurements (**Fig. 5B**). Dataset N showed particularly high ghosting and was significantly higher than all other datasets (p ≤ 0.020). Motion-equivalent temporal instability also differed substantially between rat datasets (**Fig. 5C**). Dataset B consistently showed the highest values and was significantly higher than all other rat datasets (all p < 0.001). For the mouse phantom, tSNR differed between individual datasets (**Fig. 5D**), with several significant pairwise differences but no single dataset consistently separating from all others. Ghosting showed particularly pronounced dataset-specific differences (**Fig. 5E**), with dataset R displaying substantially higher values than the remaining datasets (all p < 0.001). Motion-equivalent temporal instability also differed between datasets (**Fig. 5F**). Dataset U showed higher values than datasets E, G, H, I and R (p = 0.003 to < 0.001). Overall, the functional MRI measurements demonstrate substantial dataset-specific variability in tSNR, ghosting, and motion-equivalent temporal instability, despite acquisition at the same nominal field strength. Consistent with the findings from the anatomical measurements, the overlap across datasets indicates that field strength alone is not the sole determinant of image quality

**Figure 5:**
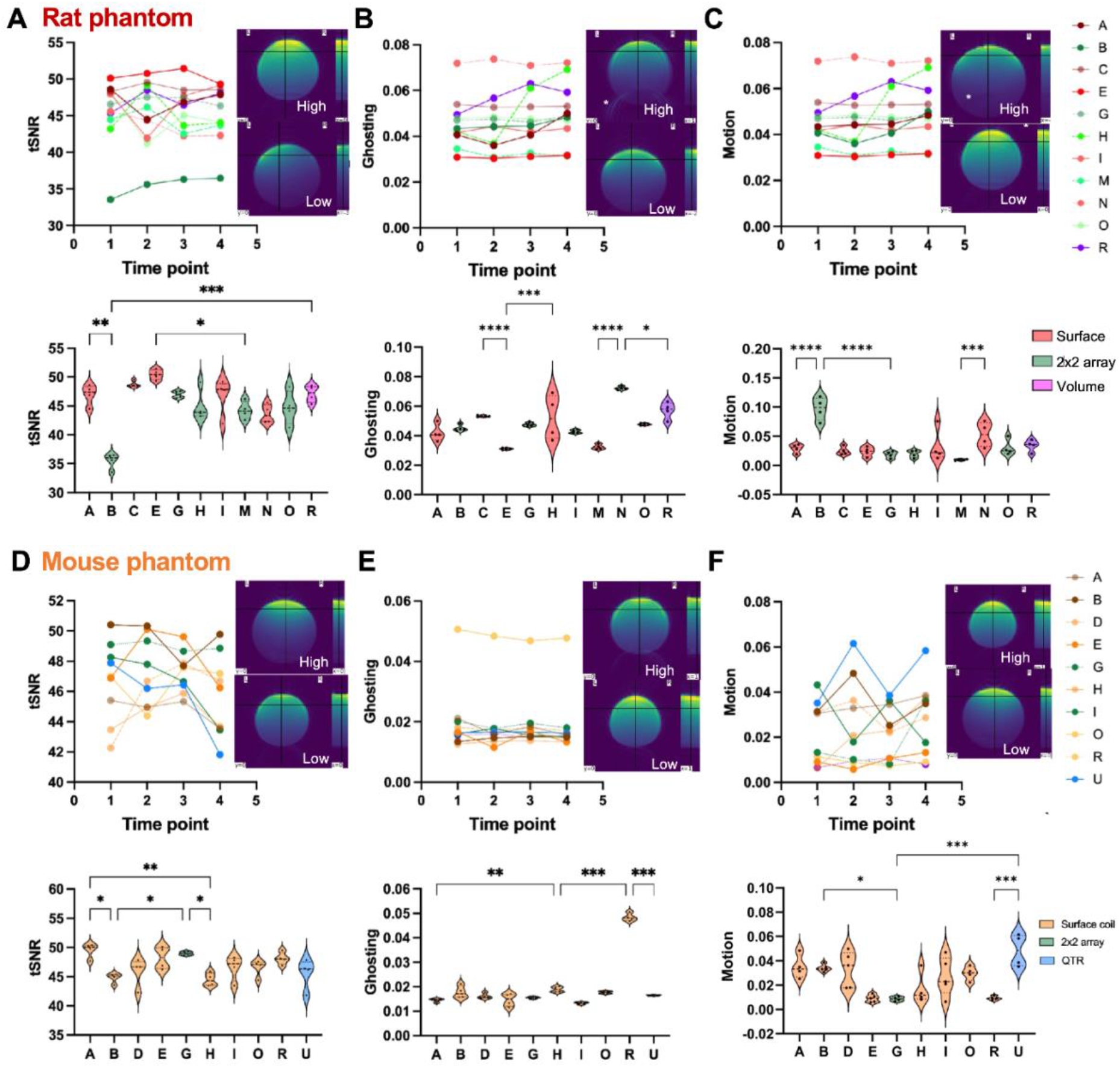
MRI quality measures for functional scans acquired with rat and mouse phantoms at 7 T. Quality control measures are shown separately for the rat (A–C) and mouse (D–F) phantoms, including temporal signal-to-noise ratio (tSNR; A, D), ghosting (B, E), and motion-equivalent temporal instability (C, F). Upper panels show individual datasets across four repeated time points; representative AIDAqc output images illustrate examples of high and low values for each quality metric (* indicates visible image artifacts). Lower panels compare the distributions of quality metric values across the four time points between datasets. Violin plots show the distribution of values, with quartiles (dotted horizontal lines) and the median (solid horizontal line). Statistics: Brown–Forsythe and Welch ANOVA followed by Dunnett’s T3 multiple-comparisons test (A) and ordinary one-way ANOVA followed by Tukey’s multiple-comparisons test (B–F). Selected significant pairwise comparisons are indicated in the figure. Detailed statistical results are available in the online data repository.

### Multicenter reference values for 7 T and longitudinal stability

For the 7 T datasets, we calculated multicenter reference values for anatomical and functional imaging across the most frequently represented configurations (**Table 1**). Specifically, we evaluated the mean SNR, tSNR, motion-equivalent temporal instability index, and ghost-to-signal ratio for mouse setups utilizing surface coils (n = 8) and rat setups utilizing 2×2 array coils (n = 7). In the mouse phantom acquired with a surface coil, the mean anatomical SNR was 48 ± 3, and the mean functional tSNR was 48 ± 1. The corresponding artifact metrics demonstrated a low predisposition to artifacts, with a mean motion of 0.03 ± 0.02 and identical ghosting of 0.02 ± 0.01 for both anatomical and functional acquisitions. For the rat phantom employing a 2×2 array coil, the established reference values yielded a mean anatomical SNR of 49 ± 3, and a functional mean tSNR of 44 ± 4, i.e., with higher variability. While the mean motion index for the rat setup remained identical to the mouse configuration at 0.03 ± 0.02, the mean ghosting level exhibited a slight increase to 0.05 ± 0.01 across both sequences. These values provide empirical multicenter references for the represented 7 T configurations rather than universal acceptance threshold

**Table 1:**
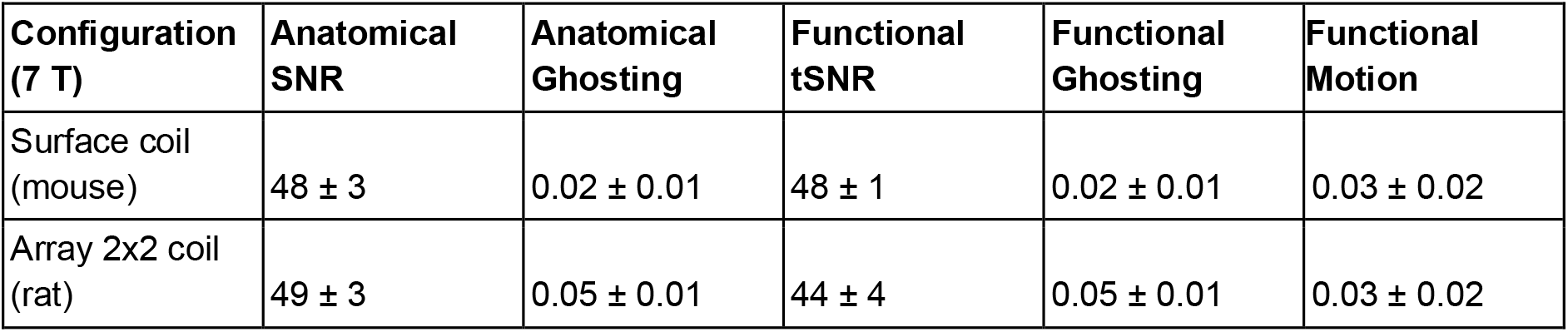
Reference quality control values for anatomical and functional MRI at 7 T. Data represent the mean and standard deviation (± SD) of quantitative quality metrics derived from standard preclinical setups: a mouse phantom acquired with a surface coil (n=8) and a rat phantom acquired with a 2×2 array coil (n = 7). Evaluated metrics include anatomical SNR, functional tSNR, ghosting levels (assessed in both sequences), and functional motion index.

| Configuration (7 T) | Anatomical SNR | Anatomical Ghosting | Functional tSNR | Functional Ghosting | Functional Motion |
| --- | --- | --- | --- | --- | --- |
| Surface coil (mouse) | $48 \pm 3$ | $0.02 \pm 0.01$ | $48 \pm 1$ | $0.02 \pm 0.01$ | $0.03 \pm 0.02$ |
| Array 2x2 coil (rat) | $49 \pm 3$ | $0.05 \pm 0.01$ | $44 \pm 4$ | $0.05 \pm 0.01$ | $0.03 \pm 0.02$ |

Temporal stability of the anatomical phantom measurements was quantified using the coefficient of variation (CV%), where lower values indicate greater temporal stability, and normalized drift (% per time point), where positive and negative values indicate increasing and decreasing values over time, respectively. Temporal-stability measures were calculated using all available longitudinal time points for each dataset. A small number of dataset-level stability measures were identified as statistical outliers and excluded from the corresponding group comparisons (**Supplementary Table 3**). Importantly, an extreme CV identifies unusually high temporal variability, whereas an extreme normalized-drift estimate does not necessarily indicate a consistent linear temporal trend. Regression-derived 95% confidence intervals (CIs) were therefore additionally considered when interpreting normalized drift.

For the rat phantom, SNR showed generally low temporal variability **(Fig. 6A).** Mean SNR CV was 1.42% at 7 T and 2.34% at 9.4 T after exclusion of dataset S at 9.4 T (CV = 18.94%); the single 3 T dataset showed a CV of 7.48% and was included descriptively. Normalized SNR drift was close to zero at 7 T and small at 9.4 T (Fig. 6B). Dataset G was an exception at 7 T, showing a significant negative SNR drift of −1.77% per time point (95% CI, −3.49 to −0.05%). Ghosting showed greater temporal variability than SNR **(Fig. 6C),** with mean CVs of 3.23% at 7 T and 6.44% at 9.4 T; dataset G was again identified as an outlier (CV = 17.90%). Mean normalized ghosting drift was close to zero at both field strengths (Fig. 6D). Although dataset G showed a large positive drift estimate, its wide 95% CI included zero, indicating pronounced variability rather than a consistent linear trend.

**Figure 6:**
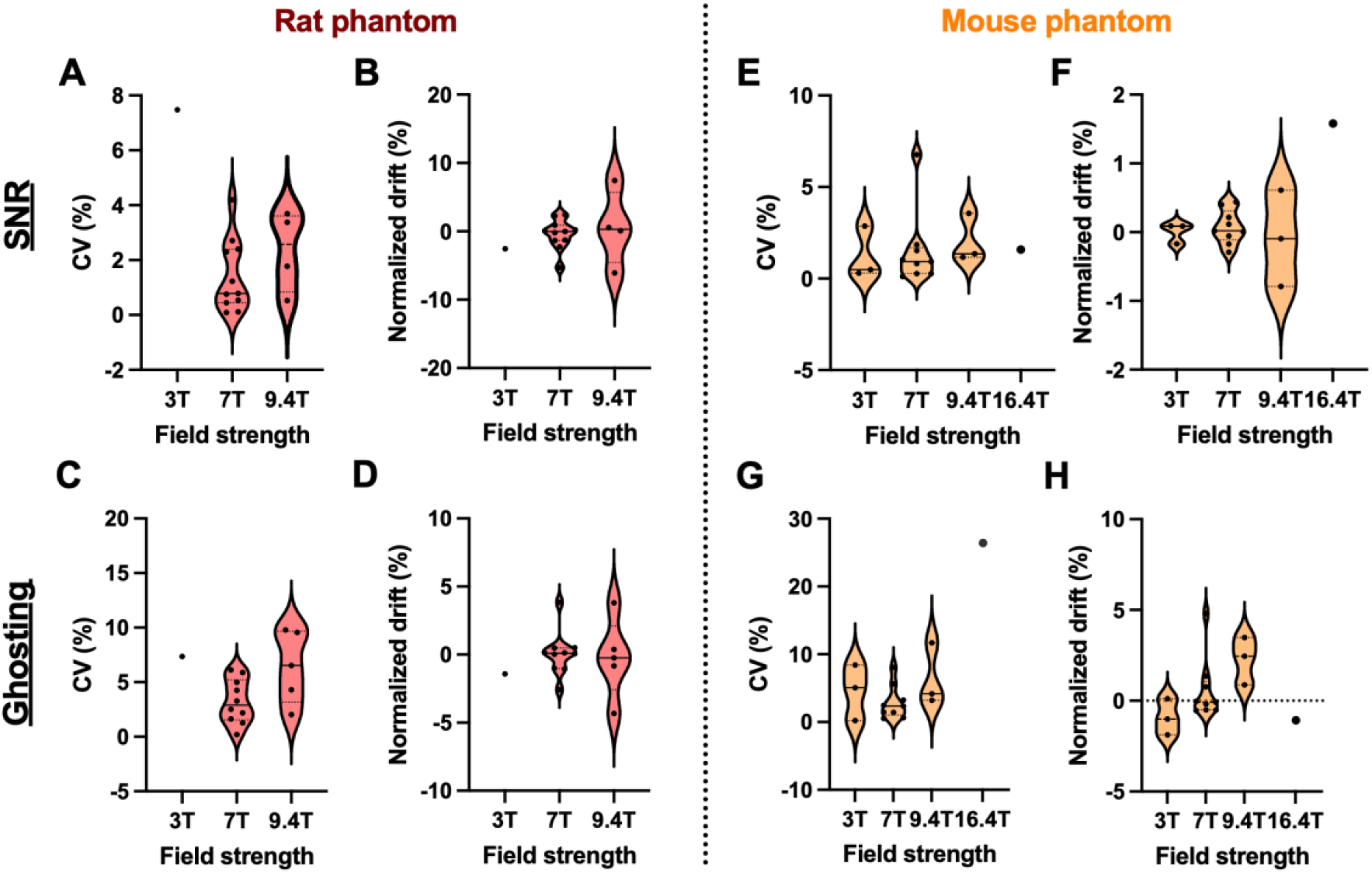
Temporal stability of SNR and ghosting for anatomical scans. Temporal stability was assessed separately for rat (A–D) and mouse (E–H) phantoms using all available longitudinal T2_RARE acquisitions for each dataset. SNR stability is shown as coefficient of variation (CV%; A, E) and normalized drift (B, F), and ghosting stability as CV% (C, G) and normalized drift (D, H). When multiple acquisitions were available for the same dataset and time point, values were averaged before calculation of temporal-stability measures. CV% was calculated for each dataset as the standard deviation across all available unique time points divided by the corresponding mean and multiplied by 100. Normalized drift was derived from the slope of a linear regression across all available time points, normalized to the dataset mean, and expressed as percentage change per time point. Positive and negative normalized-drift values indicate increasing and decreasing values over time, respectively. Each point represents one independent laboratory dataset. Data are shown as violin plots, with dotted lines indicating quartiles and solid lines the median. Rat phantom data are shown for 3 T (n = 1), 7 T (n = 11), and 9.4 T (n = 5). The single 3 T dataset is shown descriptively but was excluded from inferential statistics. Rat 7 T and 9.4 T datasets were compared using Welch’s t-test for SNR CV%, SNR normalized drift, and ghosting CV% (A–C) and the Mann–Whitney test for ghosting normalized drift (D). Mouse phantom data are shown for 3 T (n = 3), 7 T (n = 10), 9.4 T (n = 3), and 16.4 T (n = 1). The single 16.4 T dataset is shown descriptively but was excluded from inferential statistics. Differences between 3, 7, and 9.4 T were assessed using Welch’s ANOVA for SNR CV% and normalized drift (E, F) and Kruskal–Wallis tests for ghosting CV% and normalized drift (G, H). Observations identified as statistical outliers by Grubbs’ test were excluded from the corresponding group-level analyses and are reported in Supplementary Table S3. No significant differences between field strengths were detected for any temporal-stability measure (p > 0.05).

For the mouse phantom, SNR similarly showed low temporal variability across field strengths **(Fig. 6E).** After exclusion of dataset I at 7 T (CV = 13.19%), mean SNR CV ranged from 1.22% to 2.03% across 3, 7, and 9.4 T. Normalized SNR drift was close to zero across these field strengths (Fig. 6F). Ghosting was again more variable than SNR **(Fig. 6G),** with mean CVs ranging from 2.86% to 6.36% after exclusion of dataset U at 7 T (CV = 66.92%). Normalized ghosting drift showed no consistent pattern across field strengths (Fig. 6H). Dataset U had an extreme negative drift estimate, but its wide 95% CI included zero, indicating high temporal variability rather than a consistent decrease. The single 16.4 T dataset was included descriptively and showed low SNR variability (CV = 1.58%) but substantially higher ghosting variability (CV = 26.42%).

For both mouse and rat phantoms, neither temporal variability (CV%) nor normalized drift of SNR and ghosting differed significantly between the field strengths included in the inferential comparisons. Overall, SNR showed relatively low temporal variability and limited systematic drift across most datasets, whereas ghosting showed greater temporal variability and pronounced changes in individual datasets. These findings indicate that longitudinal stability was characterized primarily by dataset-specific deviations rather than systematic differences between field strengths.

### Detection of acquisition instabilities

While temporal stability was generally high across laboratories, specific acquisition instabilities were detected by the machine learning-based majority voting in a subset of the longitudinal datasets, of which representative examples (from all votings.csv files) are discussed in detail (**Fig. 7**). Here datasets up to time point 5 were included, which was not included in the previous analysis. Majority voting ranges from 0 - 5 with 1 representing one classifier, which labeled the time point as a technical outlier (**Fig. 7A**). For robust outlier detection in the mouse phantom dataset H (majority voting > 3), we identified a pronounced drop in tSNR and increase in motion at time point 5 (**Fig. 7A**). Similarly, dataset G exhibited an abrupt increase in motion and decrease in tSNR at time point 4 (**Fig. 7B)**. In both cases, visual inspection of the individual slices revealed no apparent morphological changes or artifacts. In contrast, in dataset I, the majority voting 4 was mainly driven by the intrusion of an artifact (likely originating from a phantom support or rim) which locally increased the noise floor and consequently degraded the SNR calculation (**Fig. 7C**).

**Figure 7:**
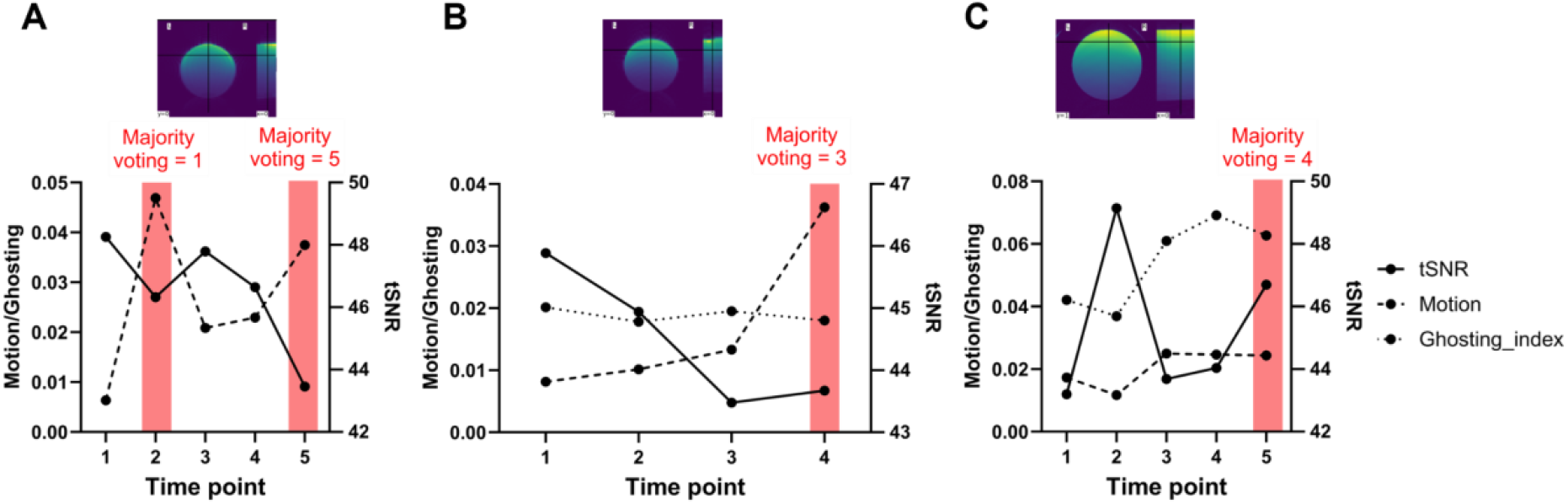
Majority voting detects temporal instabilities in longitudinal phantom quality data. Line plots illustrate deviations in tSNR (solid line), motion (dashed line) and ghosting (dotted line) across consecutive time points for functional datasets H, G, and I (A-C). Majority voting highlighting specific time points with deviating image quality (red bar) and respective phantom image (false-colored).

## Discussion

In preclinical MRI using small animal models, basic quality control protocols on how to control for scanner stability, measurement errors, and drift in longitudinal scans remain a rather manual approach with no standardized workflow, timing, and reporting. This is in stark contrast with common practice in radiology. Here, we compared longitudinal mouse and rat phantom data from 21 sites representing the heterogeneity of the field in terms of field strength (3 - 16.4 T), coil designs (loop, array, and volume design), using identical MR sequences. SNR and tSNR generally increased with field strength, whereas artifact-related measures showed more heterogeneous field- and dataset-specific patterns. At 7 T, we characterized between-dataset variability across commonly used scanner and coil configurations and provide multicenter reference values for frequently represented configurations. These values should be considered empirical references rather than universal acceptance thresholds, as substantial variability remained even among datasets using the same nominal coil configuration. Scanner- specific longitudinal baselines may therefore be more informative for detecting changes in system performance. By leveraging the standardized AIDAqc pipeline, identical and fully reproducible extraction of quality metrics (e.g., assessing the performance of EPI sequences, field strengths, and coils) was guaranteed across datasets. This workflow comprises the first small animal MRI QC/QA protocol to be established as a community-driven standard.

### Field Strength Effects

While theoretical expectations and prior single-center studies suggest that signal metrics generally increase with magnetic field strength (Edelstein et al. 1986; Pohmann et al. 2016), our multicenter data revealed a more nuanced picture. Higher SNR and tSNR were observed in the mouse phantom at 7 and 9.4 T compared with 3 T, whereas the rat phantom showed no significant differences in SNR between field-strength groups and only a non-significant increase in tSNR from 7 to 9.4 T. Motion-equivalent temporal instability and ghosting did not differ significantly between field-strength groups in either phantom. Importantly, field strength was not investigated in isolation: the groups comprised heterogeneous hardware and acquisition configurations, including different RF coils, spatial resolutions, and sequence parameters such as TR and TE. These factors likely contributed to the within-group variability and may have influenced the observed differences in signal metrics, emphasizing that nominal field strength alone is insufficient to predict image-quality performance in a multicenter setting. The single 16.4 T mouse dataset further illustrates the importance of considering signal- and artifact-related QC metrics separately. Although one dataset cannot support conclusions about ultra-high-field performance, differences among its signal, ghosting, and temporal- instability metrics demonstrate that favorable performance in one QC domain does not necessarily imply favorable performance in another. This distinction is particularly relevant for EPI-based acquisitions, which are sensitive to B0 and B1 inhomogeneities, gradient imperfections, and phase inconsistencies (Ladd et al. 2018; Obriot et al. 2026). In a multicenter context, such effects can impair comparability if they are not explicitly monitored. Our findings therefore support evaluating ghosting and temporal instability alongside SNR and tSNR rather than relying on signal metrics alone (Friedman and Glover 2006).

### Influence of RF Coil Configuration

RF coil design is an important determinant of MRI sensitivity and spatial sensitivity profiles, because coil geometry, filling factor, sample proximity, and array configuration affect SNR and signal homogeneity (Doty et al. 2007). However, at constant field strength, our 7 T data did not show a consistent separation of image-quality metrics according to nominal coil configuration. At 7 T, image-quality metrics varied substantially between individual datasets, including among datasets using the same nominal RF coil configuration. No consistent pattern across coil categories was apparent. This is expected because RF-coil performance depends on multiple factors, including coil geometry, dimensions, loading, and coil-to-sample distance (Doty et al. 2007). Similarly, the rat datasets showed substantial overlap between surface- and array-coil acquisitions for anatomical SNR and ghosting, while marked outlying values for functional tSNR, ghosting, or motion-equivalent temporal instability were associated with individual datasets rather than uniformly with one coil category. This observation is compatible with the broader literature showing that RF coil performance depends not only on coil class, but also on filling factor, coil-to-sample distance, tuning and matching, decoupling, loading, and the transmit/receive configuration (Doty et al. 2007).

These observations are particularly relevant for multicenter studies, where coil selection may co-vary with scanner platform, acquisition settings, positioning, and reconstruction. Previous multicenter phantom measurements at 7 T demonstrated substantial between-scanner variability in geometric accuracy and showed that standardized acquisition reduced variability compared with locally implemented protocols (Milidonis et al. 2016). Importantly, calibration of a poorly performing system markedly reduced geometric errors, demonstrating that nominal field strength alone does not define system performance. Our findings extend these observations beyond geometric accuracy by demonstrating dataset-specific variability in both anatomical and functional image-quality metrics, including SNR, tSNR, ghosting, and motion- equivalent temporal instability. Collectively, these results support harmonized QA/QC procedures and longitudinal system-specific reference baselines (Osborne et al. 2017) rather than assuming equivalence based on nominal field strength or RF coil category.

Collectively, our findings therefore argue against treating either scanners or nominal RF coil categories as interchangeable in preclinical MRI QA. Rather than defining universal coil-specific acceptance criteria from the present dataset, longitudinal scanner- and configuration-specific reference baselines may provide a more robust framework for identifying deviations in system performance.

### Longitudinal Stability and Detection of Hidden Instabilities

A key strength of the proposed framework is its ability to characterize longitudinal scanner behavior. Repeated anatomical phantom measurements demonstrated generally low temporal variability and limited systematic drift of SNR, whereas ghosting was more variable, including substantially larger CVs in individual datasets. Neither CV nor normalized drift differed significantly between the investigated field strengths, suggesting that longitudinal stability was primarily dataset-specific rather than systematically determined by field strength. Importantly, CV and normalized drift provided complementary information: several datasets with high variability also showed large drift estimates, but wide 95% confidence intervals indicated that these fluctuations were not necessarily consistent linear trends. Longitudinal phantom QC is therefore valuable for establishing scanner-specific baselines and distinguishing persistent system characteristics from temporal deviations.

The multialgorithm analysis additionally identified isolated acquisitions with abrupt changes in SNR, tSNR, ghosting, or temporal instability, some of which were not apparent from visual inspection alone. Such deviations may arise from scanner drift, system upgrades, hardware or calibration changes, or acquisition-related variability. Previous longitudinal MRI studies have demonstrated that scanner drift, inter-scanner variability, and even software upgrades can measurably affect quantitative imaging results, while longitudinal phantom measurements can reveal scanner instabilities that propagate into functional imaging analyses (Takao et al. 2011; Kayvanrad et al. 2021; Takao et al. 2012). Such technical changes could be misinterpreted as biological effects in longitudinal studies if appropriate QA data are unavailable.

From an operational perspective, this study provides strong evidence that routine, standardized QC is both feasible and informative in preclinical MRI, even across institutions with diverse equipment and expertise. The results support several practical recommendations (Box 1).

1. Routine phantom-based QA should be performed at regular intervals, ideally weekly depending on scanner usage and experimental sensitivity.
2. QA metrics should extend beyond basic SNR to include tSNR, ghosting, and motion-equivalent temporal instability, particularly for functional imaging workflows.
3. Reference ranges should account for field strength, coil and acquisition configuration, while longitudinal scanner-specific baselines should be used to identify deviations from established performance.
4. QA results should be reviewed longitudinally to detect trends and step-changes, not only single-time-point outliers.

Importantly, the data demonstrate that a lack of dramatic failures does not imply the absence of meaningful variability. If unrecognized, such technical variability can increase measurement variance and thereby reduce statistical power and reproducibility.

### SOPs and Harmonization Across Preclinical and Clinical Domains

Compared to clinical MRI, where QA/QC procedures are often mandated and standardized (Ihalainen et al. 2011; Chen et al. 2004), preclinical imaging continues to rely heavily on local practices and expert knowledge (Osborne et al. 2017; Tavares et al. 2023). This study highlights the consequences of this gap but also outlines a concrete path forward.

Standard Operating Procedures for preclinical MRI QA should explicitly define: 1) Phantom type and composition, 2) Acquisition protocols and parameter ranges, 3) Required metrics and reporting formats, 4) Frequency of measurements, 5) Action thresholds and escalation procedures. While full harmonization equivalent to clinical settings may not be realistic, the adoption of minimal common standards would substantially improve data interoperability and reusability (Osborne et al. 2017; Tavares et al. 2023; Briggs et al. 2021; Inau et al. 2021). The FAIR-compliant data and code sharing approach demonstrated here further enables transparent reference values across laboratories and vendors.

### Extendibility to Other Modalities

Although the present work focused on structural and EPI-based functional MRI, the framework is readily extendable to other modalities commonly used in preclinical research. Diffusion-weighted imaging, spectroscopy, and quantitative relaxometry all suffer from similar vulnerabilities to hardware drift and site-specific biases. Incorporating modality-specific phantoms and metrics into a unified QC ecosystem (Ricchi et al. 2025; Song et al. 2015; Emmerich et al. 2021) represents a logical next step toward comprehensive scanner health monitoring and protocol validation.

### Limitations and User Responsibility

Several limitations should be acknowledged. First, comparisons were not fully balanced across field strengths, coil types, acquisition settings, and time points, reflecting the realities of a community-driven multicenter study. This heterogeneity was maintained to evaluate the framework under realistic conditions. SNR was used as an image-quality QC metric and was not normalized for voxel volume or acquisition time; between-site differences should therefore not be interpreted as direct measures of intrinsic scanner sensitivity. Moreover, T1 and T2 of the CuSO₄-containing phantoms were not measured at the different field strengths, and field-dependent relaxation combined with differences in TR and TE may have contributed to the observed signal differences. Acquisition-efficiency- and relaxation-corrected SNR would enable more controlled comparisons of intrinsic scanner sensitivity but was outside the scope of the present QC framework.

Second, phantom-based QA cannot capture all sources of variability relevant to in vivo experiments, including physiological noise and animal handling. Moreover, QA tools identify variability but do not eliminate it; their effectiveness depends on informed interpretation and appropriate corrective actions. The framework should therefore be considered a decision-support rather than an automated certification system.

Phantom and FOV positioning represents an additional source of variability. Position relative to the magnet and RF-coil isocenter was not quantitatively standardized across sites, and different phantom holders were used. Position-dependent B0/B1 homogeneity, gradient nonlinearity, or holder-related artifacts may therefore affect image quality and automated ROI placement. Artifacts within the background ROIs may be interpreted as noise, resulting in an overestimation of background noise and consequently an underestimation of SNR. In addition, substantial FOV misalignment may shift the automatically defined signal or background ROIs away from their intended locations and thereby bias the SNR estimate. Standardized holders and more robust landmark detection could reduce these effects.

The implemented SNR method estimates noise from eight corner regions of a single magnitude image. Although pooling these regions reduces dependence on individual ROI placement, it cannot fully account for spatially varying or correlated noise introduced by multi-element coils, parallel imaging, or reconstruction filterings (Dietrich et al. 2007). AIDAqc SNR values should therefore be interpreted as relative QC indicators, particularly when comparing different coil or reconstruction configurations. Finally, the framework does not include a dedicated geometric phantom for detecting gradient-related scaling changes and nonlinear spatial distortions, nor does it correct systematic effects such as longitudinal gradient scaling changes or gradient nonlinearity, as implemented in established multicenter MRI QC frameworks (van Houdt et al. 2020).

## Data and code availability

The dataset including all raw and processed MRI data (https://doi.org/10.12751/g-node.s9ezrq), and code (https://github.com/Aswendt-Lab/AIDAqc) are available under GitHub and Gin repository (CC BY-NC-SA 4.0 license) according to a standardized FAIR data scheme (Kalantari et al. 2023).

## Acknowledgments

We gratefully acknowledge the technical assistance and image acquisition performed by a staff member of the France Life Imaging network (ANR-11-INBS-0006), as well as the technical support provided by Britt D’Hauw from the Bio-Imaging Lab (Antwerp, Belgium). The Italian Ministry for Education and Research (MIUR) is acknowledged for its annual FOE funding to the Euro-BioImaging Multi-Modal Molecular Imaging Italian Node (MMMI). We also thank the Queensland NMR Network and the National Imaging Facility (a National Collaborative Research Infrastructure Strategy capability) for supporting the operation of the 16.4 T MRI at the Centre for Advanced Imaging, The University of Queensland.

## Author contribution statement

BY: Methodology, resources, data curation, visualization, formal analysis, and lead in manuscript writing. AK: Software, writing. DB (Bolster, Declan): Resources, writing. EB (Botto, Elena): Resources, writing. LC: Resources, writing. SD: Resources, writing. FG: Resources, writing. MG: Resources, writing. KH: Resources, writing. JK: Resources, writing. NK: Resources, writing. XL: Resources, writing. BM: Resources, writing. EM (Micotti, Edoardo): Resources, writing. FM: Resources, writing. SM: Resources, writing. LP: Resources, writing. CP: Resources, writing. KS: Resources, writing. AT: Resources, writing. JV: Resources, writing. AV: Resources, writing. IV: Resources, writing. PW: Resources, writing. TB: Resources, writing. DB (Bertoglio, Daniele): Resources, writing. PB: Resources, writing. EB (Budinger, Eike): Resources, writing. JD: Resources, writing. RD: Resources, writing. CF: Resources, writing. GF: Resources, writing. PG: Resources, writing. WH: Resources, writing. DL: Resources, writing. MM: Resources, writing. EM (Muñoz-Moreno, Emma): Resources, writing. ER: Resources, writing. RV: Resources, writing. GI: Conceptualization, resources, methodology, formal analysis, writing. MA: Conceptualization, methodology, software, data curation, visualization, formal analysis, writing, supervision, funding acquisition.

All authors read and approved the final manuscript.

## Footnotes

1 https://github.com/Aswendt-Lab/MRI_Standardization_AIDAqc

