## Supplementary Material for "A harmonized phantom MRI quality control framework identifies sources of longitudinal and multi-site variability"

**Table S1. Summary of MRI acquisition parameters across sites.** Asterisks (*) denote minor protocol adaptations that were allowed by local hardware constraints or system specifications.

| Magnetic Field Strength (T) | Se-quence | TR/TE (ms) | RARE Factor | Band-width (KHz) | Av-erages | Repetitions | Fat sup-pressio n | Auto-ghost cor-rection |
|---|---|---|---|---|---|---|---|---|
| 3 | T1 RARE | 700/12 | 8 | 20.83 | 3 | 1 | On | X |
| 3 | T2 RARE | 2000/12 | 8 | 32.36 | 8 | 1 | On | X |
| 3 | GE EPI | 1000/12 | X | 300 | 1 | 600 | On | On |
| 7 | T1 RARE | 750/6 | 8 | 87.72 | 3 | 1 | On | X |
| 7 | T2 RARE | 2500/33 | 8 | 54.64 | 8 | 1 | On | X |
| 7 | GE EPI | 1000/20 | X | 300 | 1 | 600 | On | On |
| 9.4 | T1 RARE | 800/6 * | 8 | 96.15 | 3 | 1 | On | On |
| 9.4 | T2 RARE | 2500/8 * | 8 | 52.08 | 8 | 1 | On | X |
| 9.4 | GE EPI | 1000/8 * | X | 300 | 1 | 600 | On | On |
| 16.4 | T1 RARE | 700/6 | 8 | 11.03 | 8 | 1 | On | X |
| 16.4 | T2 RARE | 2000/20 | 8 | 55.5 | 8 | 1 | On | X |
| 16.4 | GE EPI | 1000/6.5 | X | 400 | 1 | 600 | On | On |

**Table S2. Overview of scanner models, field strengths, and coil configurations across participants.**

| Dataset | Scanner | Field strength (T) | Phantom | Coil combination |
|---|---|---|---|---|
| A | Bruker Biospec 70/20 | 7 | Mouse | RF RES 300 1H 112/086 QSN TO AD / RF + SUC 300 1H M.BR QSN RO AD |
| A | Bruker Biospec 70/20 | 7 | Rat | RF RES 300 1H 112/086 QSN TO AD / RF + SUC 300 1H R.BR QSN RO AD |
| B | Bruker Biospec 70/20 | 7 | Mouse | RF RES 300 1H 089/072 QUAD TO AD + RF ARR 300 1H M.BR LIN RO AD AV |
| B | Bruker Biospec 70/20 | 7 | Rat | RF RES 300 1H 089/072 QUAD TO AD + RF ARR 300 1H R.BR 2x2 LIN RO AD AV |
| C | Bruker BioSpec 70/20 | 7 | Rat | Bruker 86mm VolRes + RatBrainSUC |
| D | Pharmascan 70/16 | 7 | Mouse | 1H 089/072 QSN TR / 1H M.BR. QSN |
| E | Pharmascan 70/16 | 7 | Mouse | RF RES 300 1H 112/086 QSN TO AD / RF + SUC 300 1H M.BR QSN RO AD |
| E | Pharmascan 70/16 | 7 | Rat | RF RES 300 1H 112/086 QSN TO AD / RF + SUC 300 1H R.BR QSN RO AD |
| F | Bruker Biospec Maxwell 94/17 | 9,4 | Rat | RF RES 400 1H 103/082 QSN TR AD SL + RF CP MT2 400 1H 82 R.BR1 ARR[2X2] S RO |
| G | Bruker Biospec70/30 | 7 | Mouse | RF RES 300 1H 112/086 QSN TO AD / RF + SUC 300 1H M.BR ARR. 2X2 |
| G | Bruker Biospec70/30 | 7 | Rat | RF RES 300 1H 112/086 QSN TO AD / RF + SUC 300 1H R.BR ARR. 2X2 |
| H | Bruker Biospec 70/20 | 7 | Mouse | RF RES 300 1H 112/086 QSN TO AD / RF + SUC 300 1H M.BR QSN RO AD |
| H | Bruker Biospec 70/20 | 7 | Rat | RF RES 300 1H 112/086 QSN TO AD / RF ARR 300 1H R.BR 2X2 LIN RO AD |
| I | Bruker BioSpin 70/30 | 7 | Mouse | Mouse Surface Brain RF SUC 300 1H M.BR QSN RO AD + Linear Transmitter (Surface) RF RES 300 1H 112/072 LIN TO AD |
| I | Bruker BioSpin 70/30 | 7 | Rat | Rat Surface Brain RF ARR 300 1H R.BR 2x2 LIN RO AD + Linear Transmitter (Surface) RF RES 300 1H 112/072 LIN TO AD |
| J | Bruker Biospec 30/18 | 3 | Mouse | RF RES 128 1H 103/082 QSN TR AD + RF SUC 128 1H M.BR. QSN RO AD AUTOPAC |
| J | Bruker Biospec 30/18 | 3 | Mouse | RF RES 128 1H 064/023 QSN TR |
| J | Bruker Biospec 30/18 | 3 | Rat | RF RES 128 1H 103/082 QSN TR AD + RF ARR 128 1H R.BR 2x2 RO AD |
| K | Bruker Biospec 94/16 | 9,4 | Rat | RF RES 400 1H 103/082 QSN TR + RF ARR 400 1H R.BR LIN RO AD |

| | | | | |
|---|---|---|---|---|
| K | Bruker Biospec 94/16 | 9,4 | Mouse | RF RES 400 1H 103/082 QSN TR + RF SUC 400 1H M.BR LIN RO AD |
| L | Bruker Biospec 94/16 | 9,4 | Mouse | RF RES 400 1H 112/086 QSN TO AD + RF ARR 400 1H M.BR. 2x2 RO AD |
| L | Bruker Biospec 94/16 | 9,4 | Rat | RF RES 400 1H 112/086 QSN TO AD + RF ARR 400 1H R.BR LIN RO AD |
| M | Bruker Biospec 70/20 | 7 | Rat | RF RES 300 1H 112/086 QSN TO AD + RF ARR 300 1H R.BR 2X2 LIN RO AD |
| N | Bruker PHS70/16 | 7 | Rat | RF RES 300 1H 089/072 QUAD TO AD / RF ARR 300 1H R.BR 2X2 LIN RO AD |
| O | Biospec 70/30 | 7 | Mouse | volume 72 mm + mouse quad surf coil |
| O | Biospec 70/30 | 7 | Rat | volume 72 mm + rat 2x2 surface coil |
| P | Bruker Biospec 16.4/11 | 16,4 | Mouse | 5 mm Surface Acoustic Wave animal microimaging coil |
| Q | Bruker Biospec Maxwell 30/17 | 3 | Mouse | RF RES 128 1H 103/082 QSN TR AD + RF SUC 128 1H M.BR. QSN RO AD AUTOPAC |
| R | Bruker Biospec 70/20 | 7 | Mouse | RF RES 300 1H 112/086 QSN TO AD / RF + SUC 300 1H M.BR QSN RO AD |
| R | Bruker Biospec 70/20 | 7 | Rat | RES 300 1H 075/040 QSN TR (BMRIDE T13161V3/0088) |
| S | Varian | 9,4 | Mouse | Millipede coil |
| S | Varian | 9,4 | Rat | Helmholtz + surface coil |
| T | Bruker Biospec Avance III | 9,4 | Rat | 112/086 1H transmit-receive volume resonator + 3x1 1H array coil rat head |
| T | Bruker Biospec Avance III | 9,4 | Rat | 112/086 1H transmit-receive volume resonator + 3x1 1H array coil mouse head |
| U | Bruker 300 MHz vertical magnet | 7 | Mouse | MICWB40 RES 300 1H M. BR. QTR |

**Table S3. Observations identified as statistical outliers in the temporal-stability analysis using Grubbs' test.**

| Data-set | Field strength (T) | Phantom | QC metric | Stability measure | Outlier value | 95% CI of drift |
|---|---|---|---|---|---|---|
| **Q** | 3 | Mouse | SNR | Normalized drift | −0.170%/ TP | −0.666 to 0.325 |
| **I** | 7 | Mouse | SNR | CV | 13.19% | — |
| **I** | 7 | Mouse | SNR | Normalized drift | −2.853%/ TP | −17.259 to 11.553 |
| **U** | 7 | Mouse | Ghosting | CV | 66.92% | — |
| **U** | 7 | Mouse | Ghosting | Normalized drift | −29.816%/TP | −158.829 to 99.198 |
| **G** | 7 | Rat | SNR | Normalized drift | −1.770%/ TP | −3.490 to −0.049 |
| **G** | 7 | Rat | Ghosting | CV | 17.90% | — |
| **G** | 7 | Rat | Ghosting | Normalized drift | 10.112%/ TP | −18.739 to 38.963 |
| **S** | 9.4 | Rat | SNR | CV | 18.94% | — |
